# The *Plasmodium falciparum* protein VCAP1 controls Maurer’s cleft morphology, knob architecture and *Pf*EMP1 trafficking

**DOI:** 10.1101/741033

**Authors:** Emma McHugh, Olivia Carmo, Adam Blanch, Oliver Looker, Boyin Liu, Snigdha Tiash, Dean Andrew, Steven Batinovic, Andy Low J.Y, Paul McMillan, Leann Tilley, Matthew W.A Dixon

**Author notes:** Address correspondence to Matthew W.A. Dixon. E.M. and O.C. contributed equally to this work. L.T. and M.W.A.D. contributed equally to the supervision of this work.

## Abstract

The malaria parasite, *Plasmodium falciparum*, traffics the virulence protein, *P. falciparum* erythrocyte membrane protein 1 (*Pf*EMP1) to the surface of infected red blood cells (RBCs) via membranous organelles, known as the Maurer’s clefts. We developed a method for efficient enrichment of Maurer’s clefts and profiled the protein composition of this trafficking organelle. We identified 13 previously uncharacterised or poorly characterised Maurer’s cleft proteins. We generated transfectants expressing GFP-fusions of 7 proteins and confirmed their Maurer’s cleft location. Using co-immunoprecipitation and mass spectrometry we have generated a protein interaction map of proteins at the Maurer’s clefts. We identified two key clusters that may function in the loading and unloading of *Pf*EMP1 into and out of the Maurer’s clefts. We focus on a putative *Pf*EMP1 loading complex that includes the newly characterised virulence complex assembly protein 1 (VCAP1). Disruption of VCAP1 causes Maurer’s cleft fragmentation, aberrant knobs, ablation of *Pf*EMP1 surface expression and loss of the *Pf*EMP1 directed adhesion. ΔVCAP1 parasite lines have a growth advantage compared to wildtype parasites; and the infected RBCs are more deformable and more osmotically fragile.

**Importance:** The trafficking of the virulence antigen *Pf*EMP1 and its presentation at the knob structures at the surface of parasite infected RBCs is central to severe adhesion related pathologies such as cerebral and placental malaria. This work adds to our understanding of how PfEMP1 is trafficked to the RBC membrane by defining the protein-protein interaction networks that function at the Maurer’s clefts controlling PfEMP1 loading and unloading. This work adds significantly to our understanding of virulence protein trafficking and will provide crucial knowledge that will be required to determine the mechanisms underpinning parasite driven host cell remodelling, parasite survival within the host and virulence mechanisms.

## Introduction

Each year, *Plasmodium falciparum* causes ∼200 million cases of illness in humans and more than 400,000 deaths, mostly of children under the age of five (1). In the asexual blood stage of infection, parasites invade red blood cells (RBCs) and develop through the so-called ring, trophozoite (growing) and schizont (dividing) stages, eventually releasing invasive merozoites that continue the blood cycle. During this cycle the parasite induces marked changes to the host RBC, including the elaboration of new organelles in the RBC cytoplasm, known as the Maurer’s clefts and the establishment of protrusions at the RBC membrane, known as knobs. The knobs comprise a spiral protein structure supported by the knob-associated histidine-rich protein, (KAHRP) (2). The knob acts as a scaffold for the presentation of the major virulence antigen *P. falciparum* erythrocyte membrane protein 1 (*Pf*EMP1). This virulence complex has an important role in adhesion of infected RBCs to endothelial cell receptors. Rigid mature stage-infected RBCs sequester away from the circulation, thus avoiding recognition and removal during passage through the spleen (3). *Pf*EMP1 is exported into the host RBC cytoplasm via a translocon at the parasitophorous vacuole membrane (4, 5). The current model suggests that *Pf*EMP1 is trafficked to Maurer’s clefts as a soluble, chaperoned complex (6, 7), where it is inserted into the membrane bilayer (8), before repackaging for vesicle-mediated trafficking to the RBC surface (9).

Maurer’s clefts are roughly disc-shaped cisternae, with a diameter of ∼500 nm (10). In the ring stage (up to ∼20 h post-invasion), they are mobile in the RBC cytoplasm but later become tethered to the RBC membrane skeleton (11). Remodelled host actin filaments and parasite-derived tethers connect the Maurer’s clefts to the RBC membrane (12, 13). However, the precise role of the Maurer’s clefts in protein trafficking remains unclear, and their composition is only partly defined (14). In particular, there is very limited information about interactions between proteins of the Maurer’s clefts and with their virulence-associated cargo (15, 16).

In this work we developed a method for enriching “mobile” Maurer’s clefts from parasites at 14-18 h post-invasion. We provide a detailed analysis of the Maurer’s cleft proteome, identifying and validating several novel resident proteins. Using super-resolution optical microscopy, we define the spatial organisation of new and established Maurer’s clefts proteins. We use co-immunoprecipitation (co-IP) and mass spectrometry to describe a network of protein interactions, identifying two *Pf*EMP1-interacting complexes, one of which we propose is responsible for loading *Pf*EMP1 into the Maurer’s clefts. We show that deletion of a partially characterised Maurer’s cleft component (*PF*3D7_1301700, CBP2/GEXP07), which we have renamed the virulence complex assembly protein 1 (VCAP1), leads to defective *Pf*EMP1 trafficking, altered Maurer’s cleft architecture, aberrant knob formation and a loss of parasite adhesion to endothelial ligands. These changes are accompanied by decreased cellular rigidity, increased osmotic fragility and a marked growth advantage.

## Results

### Enrichment of Maurer’s clefts from infected RBCs

Maurer’s clefts are parasite-derived membrane-bound structures that remain mobile within the RBC cytoplasm up until ∼20 h post-invasion (Movie S1). Leveraging this biology, we developed a protocol for purifying Maurer’s clefts from parasite-infected RBCs at 14-18 h post-invasion. Fluorescence microscopy of transfectants expressing GFP-tagged ring exported protein 1 (REX1-GFP; (17)), confirms a typical Maurer’s cleft profile (Fig 1A). REX1-GFP transfectant-infected RBCs were hypotonically lysed, and the suspension was extruded through a 27-gauge needle, adjusted to isotonicity, pre-cleared, and incubated with GFP-Trap beads (Fig 1B). Fluorescence microscopy of the beads (Fig S1A) confirms the presence of GFP-containing structures with dimensions consistent with a mixture of intact (∼500 nm) and fragmented Maurer’s clefts. Fractions from the total cell lysate, the cleared input and the unbound and bound fractions were analysed by Western blot (Fig 1C; full-length blots in Fig S1B). Due to the high haemoglobin content in the total, input and unbound fractions the samples were diluted 1:10 before electrophoresis; as a result, REX1-GFP and SBP1 were not detected in these fractions. REX1-GFP and the known Maurer’s cleft protein, skeleton-binding protein 1 (SBP1;(18, 19), were highly enriched in the bound fraction, while spectrin, a component of the host cytoskeleton was not detected, consistent with a high level of enrichment. The parasitophorous vacuole membrane (PVM) protein, exported protein-1 (EXP1) and the parasite cytoplasmic protein, glyceraldehyde-3 phosphate dehydrogenase (*Pf*GAPDH), were also absent (Fig 1C).

**Fig 1.**
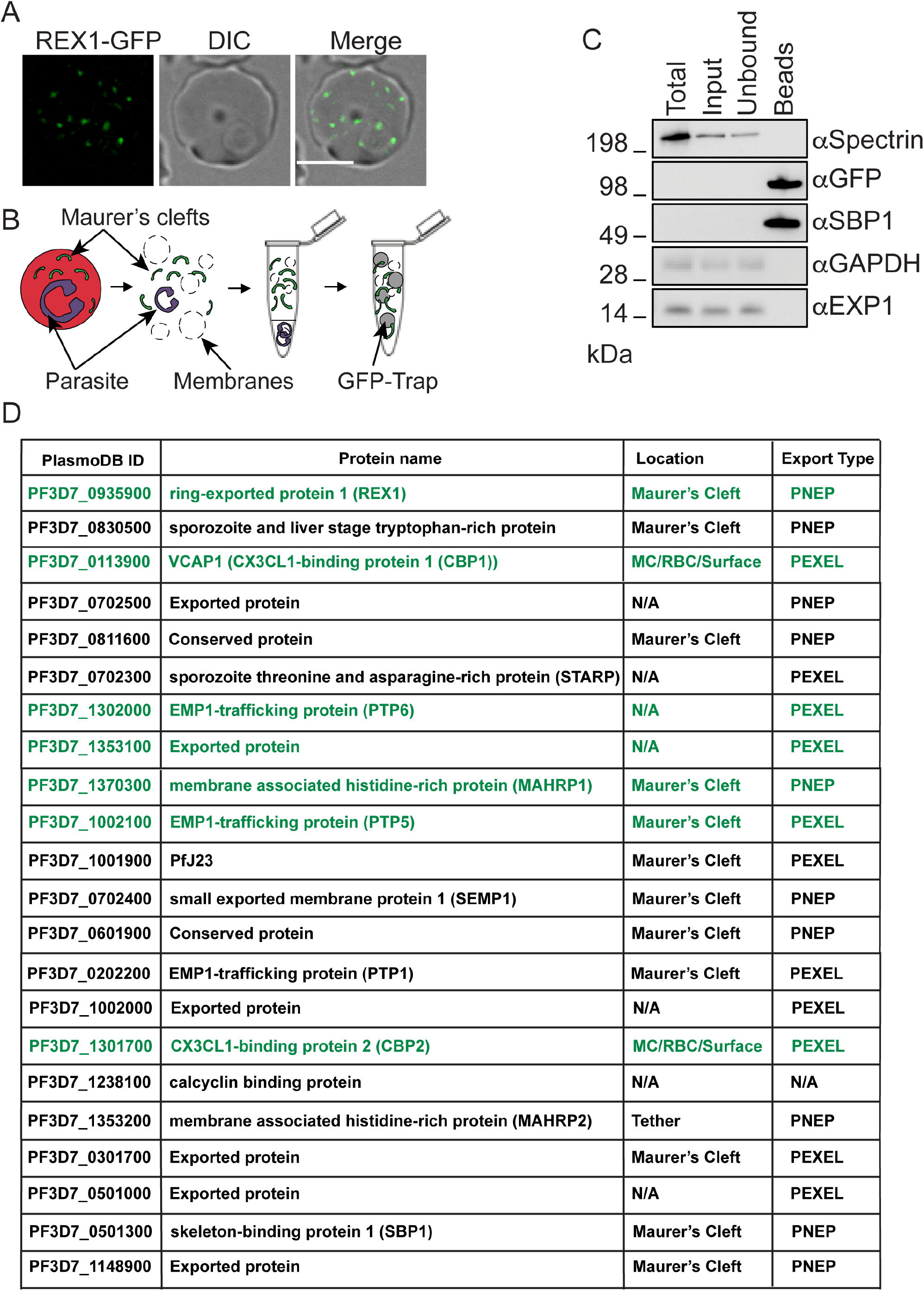
Enrichment and analysis of Maurer’s clefts from *P. falciparum* REX1-GFP infected RBCs. (A) Live cell fluorescence of an RBC infected with a REX1-GFP expressing parasite at 14-18 h post-invasion. GFP (green), DIC and merged images are shown. Scale bar 5 μm. (B) Schematic overview of the GFP-Trap Maurer’s cleft enrichment method. Infected RBCs (14 – 18 h post-invasion) are hypotonically lysed, subjected to differential centrifugation, then the REX1-GFP-labelled Maurer’s clefts are captured with GFP-Trap beads. (C) Western blot assessing enrichment of Maurer’s clefts from REX1-GFP infected RBCs. Total = complete infected RBC lysate; Input = fraction applied to the GFP-Trap beads; Unbound = supernatant after GFP-Trap bead incubation; Beads = material bound to beads. Molecular masses are in kDa. (D) Summary of proteins identified in the mass spectrometric analysis of enriched REX1-GFP Maurer’s clefts. See Data set 1 for more detail. Proteins chosen for GFP tagging are indicated in green text. MC = Maurer’s clefts; RBC = red blood cell; Surface = red blood cell surface; N/A = Not Available; PEXEL = Plasmodium export element; PNEP = PEXEL negative exported protein.

### Mass spectrometry analysis of the enriched Maurer’s clefts

The captured Maurer’s clefts were released from the beads, digested with trypsin and analysed by LC-MS/MS. Proteins were deemed significant if two or more peptides were detected in the Maurer’s cleft (REX1-GFP) sample and were 3-fold enriched compared to the 3D7 control, in 2 separate experiments (Fig 1D; Data set 1). Five well-characterised Maurer’s cleft proteins were identified, namely REX1 (20), *Pf*EMP1 trafficking protein 1 (PTP1 (21)), skeleton binding protein 1 (SBP1 (22)), small exported membrane protein 1 (SEMP1 (15)) and membrane-associated histidine-rich protein 1 (MAHRP1 (23)). A tether protein, membrane-associated histidine-rich protein 2 (MAHRP2, (13)) was also identified.

Six partially characterised proteins that have been reported previously to be Maurer’s cleft-located were identified, namely the parasite-infected erythrocyte surface protein 2 (PIESP2 (24), sporozoite and liver stage tryptophan-rich protein (25), *Pf*J23 (14), *PF*3D7_0301700, *PF*3D7_1148900, *PF*3D7_0702500 and *PF*3D7_0601900 (15, 25, 26).

We identified *PF*3D7_1301700 and *PF*3D7_0113900, for which there are conflicting reports with respect to the cellular location, ranging from the RBC cytosol in gametocytes (27), to the Maurer’s clefts in asexual stages (28), to the RBC surface (29). These proteins have been previously referred to as gametocyte exported protein 7 (GEXP07) / CX3CL1-binding protein 2 (CBP2) and GEXP10/ CBP1 (27, 29). Based on functional data described below we now refer to *PF*3D7_1301700 as the virulence complex assembly protein 1 (VCAP1).

A further 7 proteins were identified for which no location data has been reported previously. All but one protein (calcyclin-binding protein) has a predicted secretory signal or transmembrane segment and 5 have a Plasmodium export element (PEXEL) motif, which predicts export of parasite proteins to the host RBC (30). Four of these proteins have been successfully genetically disrupted: PTP5, PTP6, *PF*3D7_1353100 and *PF*3D7_0501000 (21). The remaining proteins *PF*3D7_0702300, *PF*3D7_0811600, *PF*3D7_1002000, *PF*3D7_1035800, and sporozoite threonine and asparagine-rich protein (STARP) do not have characterised locations. Over 100 peptides were observed for *Pf*EMP1 itself (Data set 1), consistent with a number of studies showing that *Pf*EMP1 is highly enriched at the Maurer’s clefts (8, 31, 32), with only a sub-population reaching the RBC membrane. Some RBC proteins were also identified in the enriched Maurer’s clefts, including Annexin A4 and A11 and Copine-3, which are calcium-dependent phospholipid-binding protein involved in membrane remodelling (33), as well as VPS28, TSG101, syntaxin and VAC14 - proteins that are involved in membrane binding, vesicle trafficking and lipid biogenesis (Table S1; Data set 1).

### GFP-tagging of proteins confirms their Maurer’s cleft location

To determine or confirm the locations of a number of the identified proteins, we generated transfectants expressing GFP fusions of five uncharacterised or partially characterised proteins (VCAP1, GEXP10/CBP1, PTP5, PTP6 and *PF*3D7_1353100) and two proteins that were previously reported to be Maurer’s cleft-located (REX1 (17) and MAHRP1), each under the control of the *CRT* (chloroquine resistance transporter) promoter (Fig 2). An immunoblot probed with anti-GFP confirms the expression of chimeric proteins that migrate close to the calculated molecular masses (Fig S1C). Using live-cell microscopy, we showed that each of the GFP-tagged proteins localises to puncta in the host RBC cytoplasm, consistent with Maurer’s cleft association (Fig 2). Immunofluorescence microscopy of samples co-labelled with antibodies recognising GFP and REX1 confirmed that the proteins are directed to the Maurer’s clefts (Fig S2A). A previous study suggested that VCAP1 and GEXP10 are present at the RBC surface (29). While the presence of a small sub-population at the RBC surface or in the RBC cytoplasm cannot be ruled out, our immunofluorescence microscopy shows that the bulk of the protein is located at the Maurer’s clefts (Fig 2; Fig S2A).

**Fig 2.**
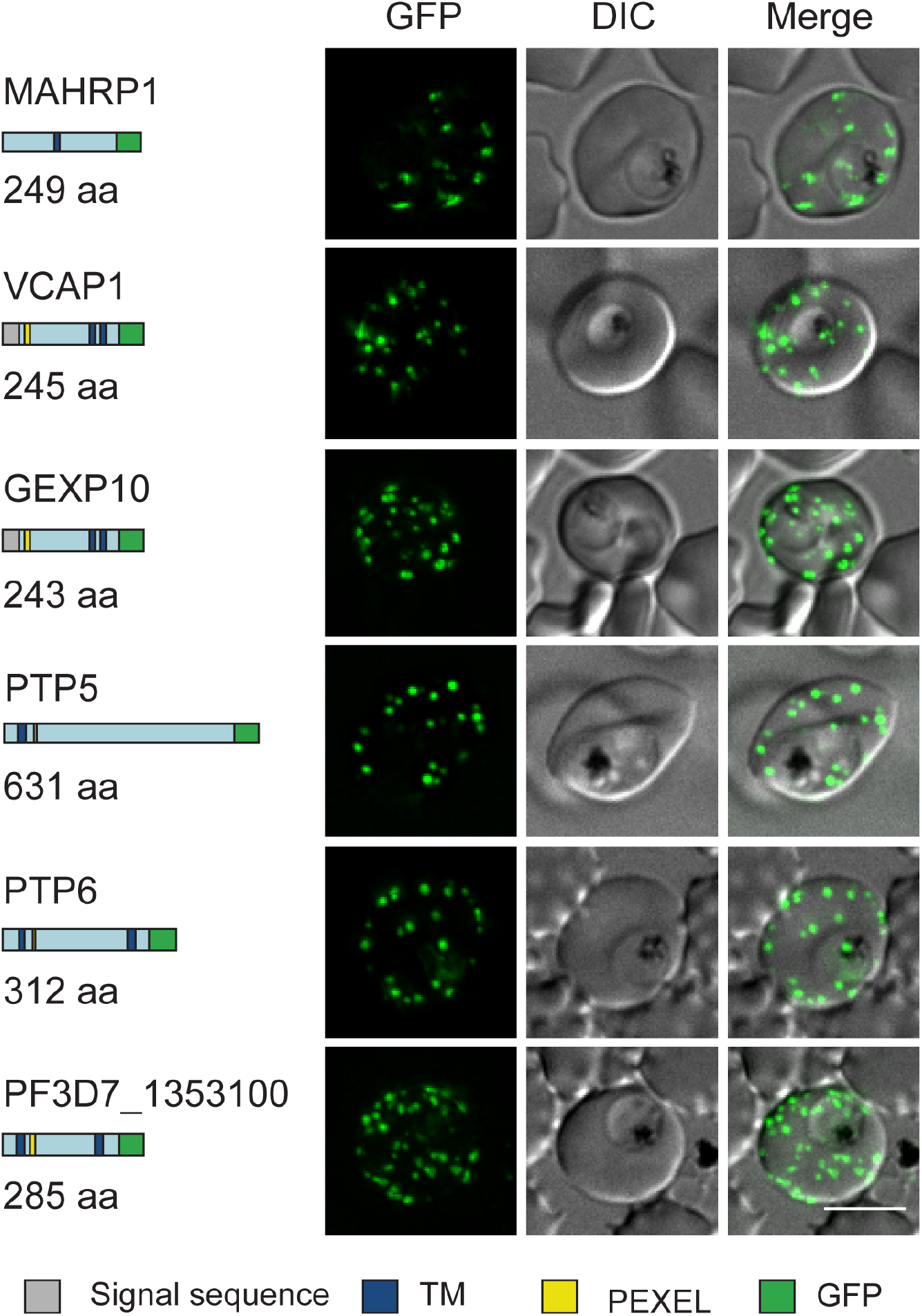
Live cell fluorescence analysis of GFP-tagged exported proteins. Predicted native protein length (amino acids) and schematic representations of six proteins that were selected for GFP-tagging are shown on the left. Grey = signal sequence, blue = transmembrane domain, yellow = PEXEL motif, green = GFP tag. Live cell fluorescence and DIC microscopy of transfectants expressing GFP-tagged putative Maurer’s cleft proteins reveals fluorescent puncta in the RBC cytoplasm. Scale bar = 5 μm.

### Co-IP of Maurer’s cleft proteins reveals a compartmentalised network of protein interaction clusters

We performed co-IP experiments on our GFP-tagged transfectants. Western analysis confirmed the enrichment of the GFP-tagged bait proteins (Fig S2B-D). In each of the lines analysed the bait protein and several interacting proteins were identified (Table S2-8; Data sets 2-8). The protein interaction networks were analysed using Navigator (Fig 3A,B, S3) (34). When combined with data from other studies that performed IPs with epitope-tagged PTP1, SEMP1 and *Pf*EMP1, a complex map of interactions is revealed (Fig S3B) (15, 16, 31). We identified 2 protein clusters that each contains Maurer’s cleft proteins that are known to have roles in *Pf*EMP1 trafficking, connecting to uncharacterised proteins and to *Pf*EMP1 itself (Fig 3A, B).

**Fig 3.**
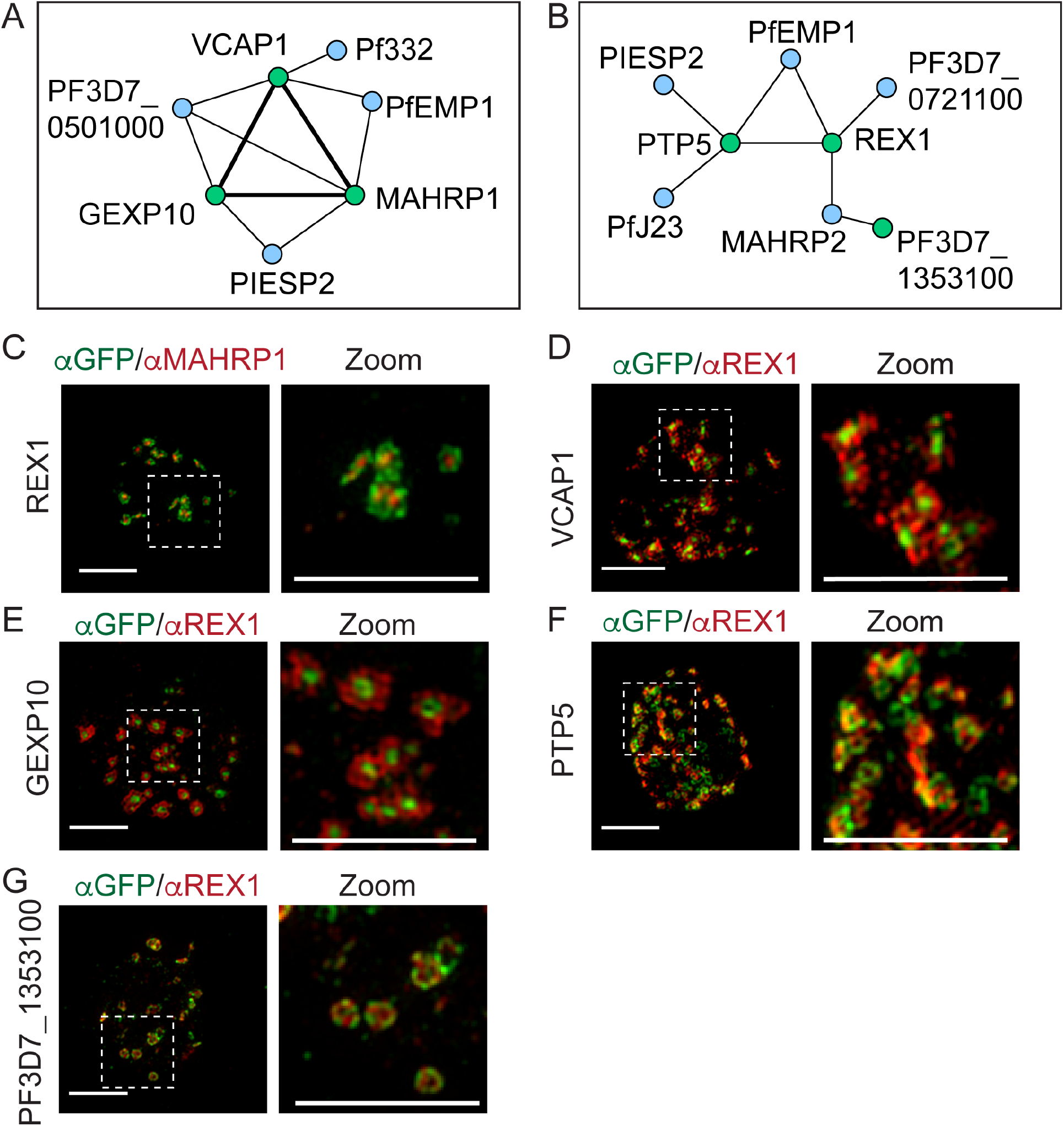
Maurer’s clefts proteins interact to form distinct protein clusters. (A) Protein interaction maps highlighting a putative *Pf*EMP1 loading hub comprising VCAP1, MAHRP1 and GEXP10 and (B) a putative unloading hub comprising REX1, PTP5, SEMP1 and *PF*3D7_1353100. Green nodes are the GFP-tagged proteins. Double thickness edges indicate reciprocal co-precipitation (See Fig S3 for full network maps). (C) 3D-SIM analysis of REX1-GFP-infected RBCs fixed and labelled with anti-GFP (green) and anti-MAHRP1 (red). (D-G) 3D-SIM analysis of transfectant-infected RBCs expressing GFP-tagged VCAP1, GEXP10, PTP5 and *Pf*3D7_1353100 that were fixed and labelled with anti-GFP (green) and anti-REX (red) antibodies. Maximum projections of Z-stacks are displayed. Scale bars = 3 μm, zoom scale bar = 3 μm.

For example, previous work showed that MAHRP1 is needed for efficient loading of *Pf*EMP1 into the Maurer’s clefts (35). We found that MAHRP1-GFP co-precipitated VCAP1 and GEXP10, while VCAP1-GFP and GEXP10-GFP co-precipitated MAHRP1 (Table S2-4; Data sets 2-4). These interactions were confirmed by Western blotting (Fig S2E). GEXP10-GFP and MAHRP1-GFP also precipitated PIESP2, while VCAP1-GFP precipitated *Pf*322, consistent with its known Maurer’s cleft location. VCAP1-GFP and GEXP10-GFP also precipitated *Pf*EMP1 (Table S2-4; Data sets 2-4), suggesting a potential role for these proteins in *Pf*EMP1 trafficking.

3D-Structured Illumination Microscopy (3D-SIM) provides an 8-fold increase in volume resolution, which enhances the analysis of compartments, such as the Maurer’s clefts, that have dimensions close to the resolution limit of conventional microscopy. 3D-SIM reveals that REX1 is located around the perimeter of the clefts, while MAHRP1 exhibits a more central location (Fig 3C, S4A), in agreement with our previous report (11). VCAP1-GFP and GEXP10-GFP are concentrated in the central region of the Maurer’s cleft surrounded by REX1 (Fig 3D,E, S4B). The profile for antibodies recognising the ATS region of *Pf*EMP1 exhibits partial overlap with that for GEXP10-GFP (Fig S4C).

The interaction analysis revealed a second protein cluster containing REX1-GFP, PTP5-GFP and *PF*3D7_1353100 (Fig 3B). REX1-GFP co-precipitated *Pf*EMP1 (Table S5; Data set 5), consistent with the known role for REX1 in trafficking *Pf*EMP1 from the Maurer’s clefts to the RBC membrane (36, 37). A co-IP with PTP5-GFP co-precipitates REX1, PIESP2 and *Pf*J23 (Table S6; Data sets 6). These data point to the existence of a protein interaction network comprised of *Pf*EMP1, REX1, PTP5 and PIESP2, potentially functioning at the cleft periphery (Fig 3B). REX1-GFP and *PF*3D7_1353100-GFP also co-precipitated the tether protein, MAHRP2 (Table S7; Data set 7).

3D-SIM imaging of PTP5-GFP and *PF*3D7_1353100-GFP cell lines reveals a dotted pattern at the periphery of the Maurer’s cleft cisternae, partially overlapping or alternating with the REX1 signal (Fig 3F, G, S4B). Similarly, *Pf*EMP1 (ATS labelling) partly overlaps with REX1 and *Pf*EMP1 at the cleft periphery (Fig S4C).

Components of the PVM translocation machinery were observed in IPs with VCAP1-GFP (HSP101, PTEX150), suggesting interactions with these proteins during the export processes. IPs with PTP6-GFP showed a number of interacting proteins including MESA, STARP and three proteins of unknown function, *PF*3D7_0702500, *PF*3D7_1002000 and *PF*3D7_0301700. This protein interaction network showed no connectivity to the clusters described above (Table S8; Data set 8).

### VCAP1 gene knockout results in altered trafficking of exported proteins and a growth advantage

As we were interested in identifying proteins that may be involved in trafficking *Pf*EMP1, we selected VCAP1 and GEXP10 (which IP each other and *Pf*EMP1) as candidates for genetic disruption. These genes have been previously reported to be refractory to deletion using conventional approaches (21, 28). In this work, we used a CRISPR-Cas9 approach to disrupt these genes in CS2 parasites (Fig S5). We were unable to obtain a knockout of GEXP10 from two separate attempts, supporting a recent genome-wide transposon screen which suggested that GEXP10 is essential (38). By contrast, a knockout was obtained for VCAP1, with disruption of the native locus confirmed by PCR (Fig S5).

We examined the ΔVCAP1 infected RBCs by immunofluorescence microscopy, using antibodies to REX1 and KAHRP (Fig 4A). An increase in the number of REX1-positive puncta was observed suggesting a change in Maurer’s cleft morphology (Fig 4A). The labelling pattern for KAHRP also appeared more punctate (Fig 4A). We next analysed the growth of the ΔVCAP1 parasite line compared with the CS2 parent parasites. The ΔVCAP1 parasite line grows faster than the parent line with a more than 2-fold difference observed after four parasite cycles (Fig 4B).

**Fig 4.**
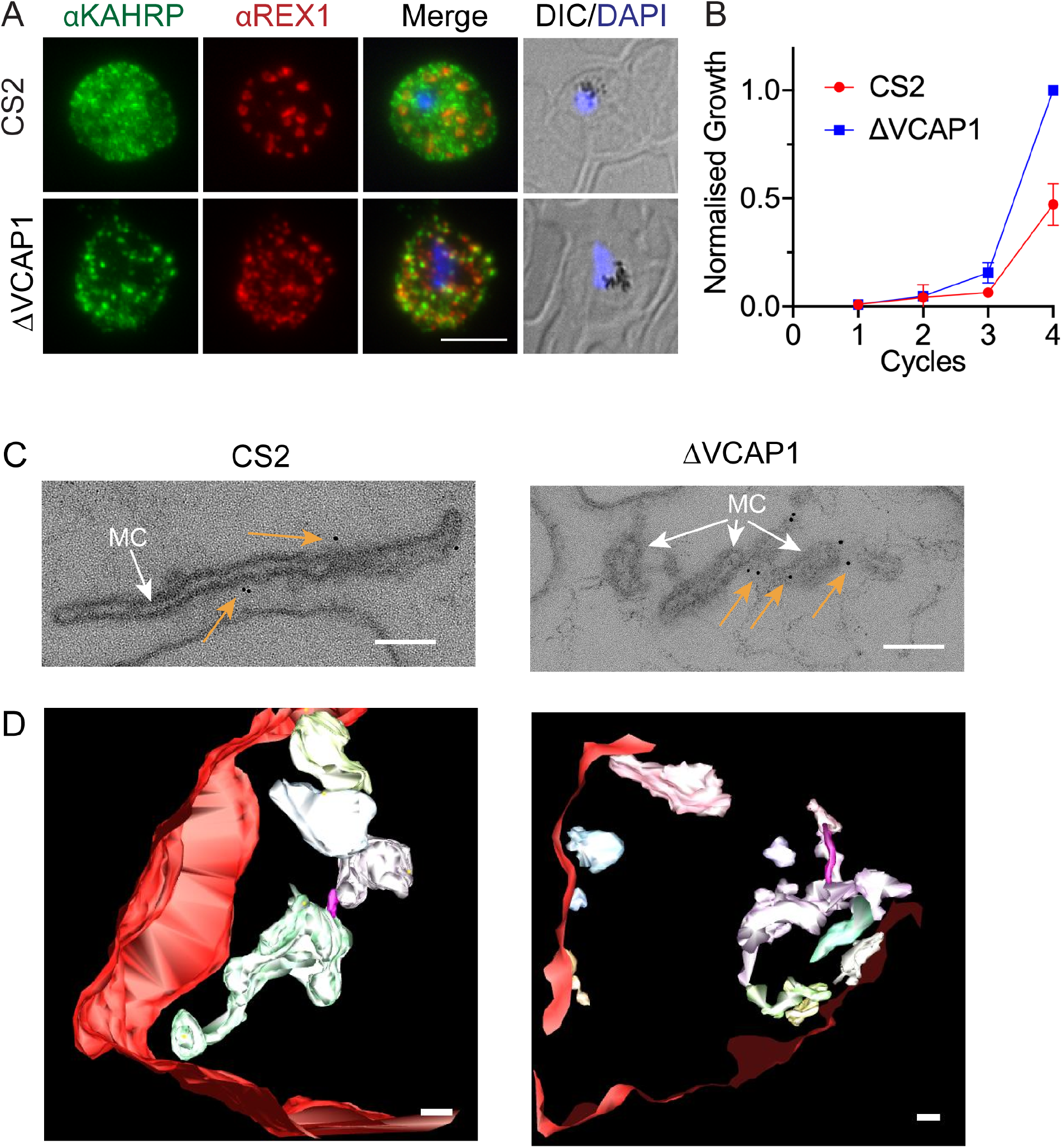
Knockout of VCAP1 alters parasite growth and Maurer’s cleft architecture. (A) Infected-RBCs were fixed and probed with anti-KAHRP, and counterstained with anti-REX1. Nuclei were stained with DAPI (blue). CS2 = parent line. Projections of Z stacks are shown. Scale bar = 5 μm. (B) Parasite growth assay measuring proliferation over 4 asexual cycles. Growth was assessed by staining infected RBCs with SYTO61 and subsequent flow cytometry analysis. The data have been normalised and expressed relative to the ΔVCAP1 parasite line (n=4). (C) EqtII-permeabilised wildtype CS2 and VCAP1 knockout transfectants were labelled with antibodies recognising REX1, followed by immunogold labelling and prepared for electron microscopy. Images have been cropped around Maurer’s clefts (white arrows) - see Fig S6A for uncropped micrographs. Gold arrows point to gold particles. Scale bar 100 nm. (D) Rendered 3D models of Maurer’s clefts generated from electron tomograms. Red = RBC; magenta stalk = tether; pastel hues = independent clefts. Scale bar = 100 nm. Translations through the virtual sections of the tomograms and rotations of the rendered models can be seen in Movies S2-3.

### Knockout of VCAP1 results in altered Maurer’s cleft morphology

RBCs infected with the ΔVCAP1 parasite line were lightly fixed and permeabilised with the pore-forming toxin, Equinatoxin II (EqtII; (39, 40)) to release haemoglobin and to permit the introduction of primary antibodies recognising REX1.

Transmission electron microscopy was performed on thin sections prepared from CS2 and ΔVCAP1 infected RBCs. The Maurer’s clefts are observed as single slender cisternae with an electron-lucent lumen and an electron-dense coat in the wild type CS2 (Fig 4C, S6A). In ΔVCAP1 infected RBCs, distorted and fragmented structures are observed (Fig 4C, S6A). These fragments label with immunogold labelled anti-REX1 antibodies, confirming a Maurer’s cleft origin (gold arrows; Fig 4C). The 3D architecture of the Maurer’s cleft fragments was examined using electron tomography (Fig 4D). The rendered images reveal the fragmented nature of the Maurer’s clefts when compared to wildtype CS2. Each separate cleft structure is rendered in a different pastel hue; the RBC (red) and the tethers (magenta) are also displayed (Fig 4D). The morphology changes are best appreciated in translations through the virtual sections of the tomogram and rotations of the rendered models (Movie S2-3).

### ΔVCAP1 parasite infected RBCs have altered knob morphology

Scanning Electron Microscopy (SEM) of intact ΔVCAP1 infected RBCs reveals enlarged knobs with a highly aberrant morphology, as well as knob clusters (Fig 5A). RBCs infected with ΔVCAP1 parasites have fewer knob structures (1.3 ± 0.2 per μm^2^) than wildtype CS2 (5.5 ± 1 per μm^2^) (Fig 5B). Thin section TEM confirmed the altered knob morphology (Fig S6B).

**Fig 5.**
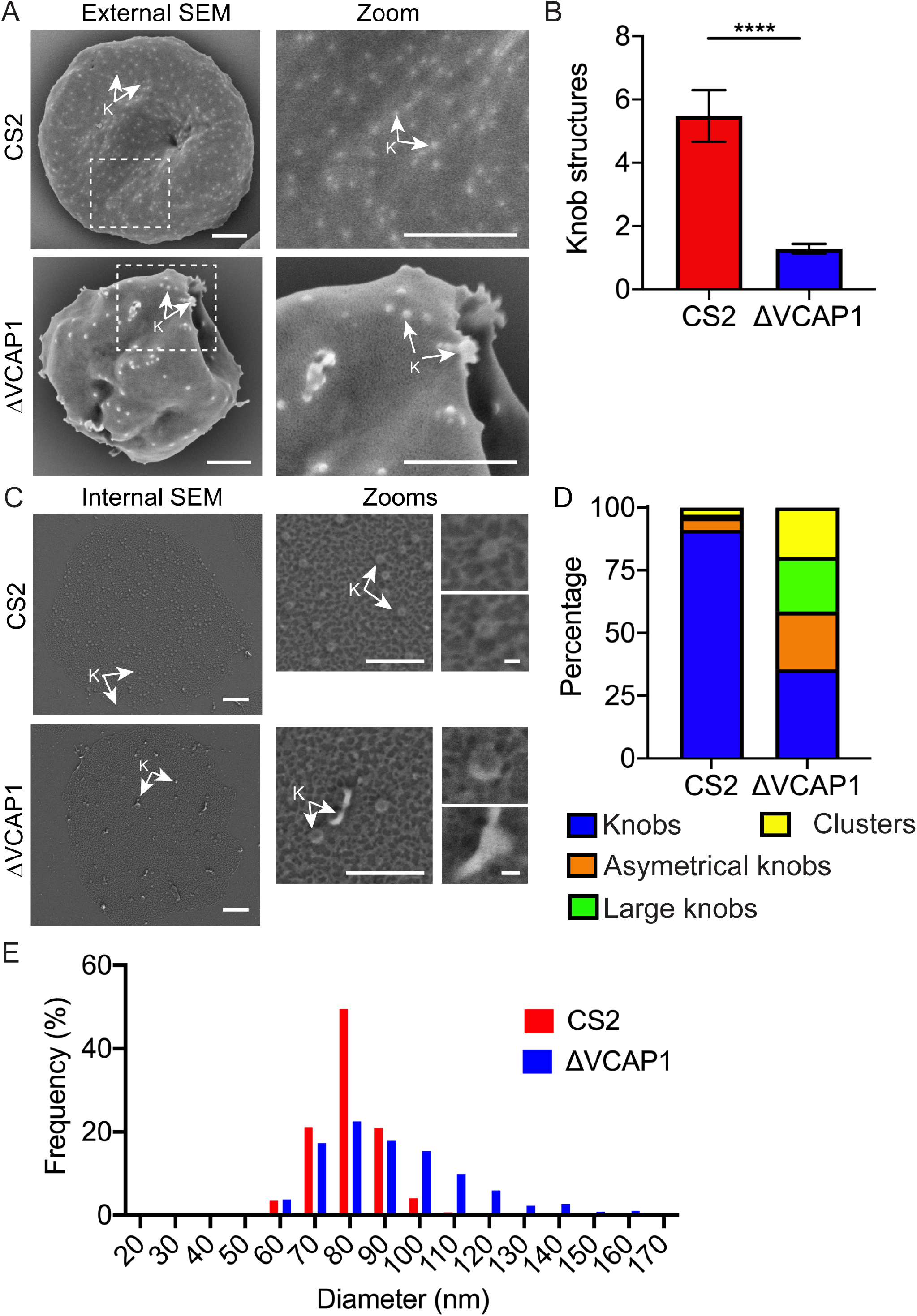
Examination of the ΔVCAP1 infected RBCs reveals altered knob morphology. (A) Mid-trophozoite stage infected-RBCs were fixed in 2.5% glutaraldehyde/PBS and prepared for SEM of the exterior surface. Scale bar = 1μm. (B) Quantification of the number of knob like structures observed in the external SEM. Data is the mean number of structures per μm^2^. (n = 15 cells per sample, minimum 3 areas per cell, Student’s t-test P<0.0001). (C) Late trophozoite stage-infected RBCs were immobilised onto glass slides. Shearing under hypotonic conditions leaves remnant membrane discs that were fixed, dehydrated, gold-coated and imaged using SEM. Knobs (K) are arrowed. Scale bar = 1μm; zoom 1 scale bar = 500 nm; zoom 2 = 50 nm. (D) Quantification of the knob morphologies observed by internal SEM imaging. These are defined as normal knobs (blue), asymmetrical knobs (orange), large knobs (green) and knob clusters (yellow). Example images of these morphologies are shown in Fig S7A. (E) Graph showing the frequency distribution of knob sizes from wildtype CS2 and ΔVCAP1 infected RBCs. A bin size of 10 nm was used; n = 760 (ΔVCAP1), 1098 (CS2).

To investigate the architecture of the knobs in more detail we made use of a recently developed method for imaging knobs at the cytoplasmic surface of infected RBCs (2). Infected RBCs were tightly linked to lectin-coated slides and subjected to shearing. The remnant membrane discs were fixed, dehydrated, gold-coated, and then imaged using high-resolution SEM. The spectrin-actin network is evident as bright skeletal elements over dark patches of background membrane (Fig 5C). The knobs appear as raised, dimpled disk-shaped structures that are closely integrated into the skeletal meshwork (Fig 5C, arrows). About 75% of the knobs were malformed in the ΔVCAP1 infected RBCs, presenting as asymmetrical, enlarged and clustered knob structures (Fig 5D, S7A). The knobs in the ΔVCAP1 infected RBC membranes exhibit a larger average diameter (90 ± 1 nm) than CS2 wild type parent (80 ± 1 nm), and a broader size distribution (Fig 5E).

### Knockout of VCAP1 ablates *Pf*EMP1 surface expression and binding to chondroitin sulphate-A

To determine if *Pf*EMP1 trafficking was affected in the ΔVCAP1 infected RBCs we performed immunofluorescence microscopy using antibodies directed towards the conserved acidic terminal segment (ATS) of *Pf*EMP1. Wildtype CS2 infected RBCs exhibit characteristic localisation of *Pf*EMP1 at the Maurer’s clefts, while *Pf*EMP1 labelling is much weaker in the ΔVCAP1 infected RBCs (Fig 6A, S7B).

**Fig 6.**
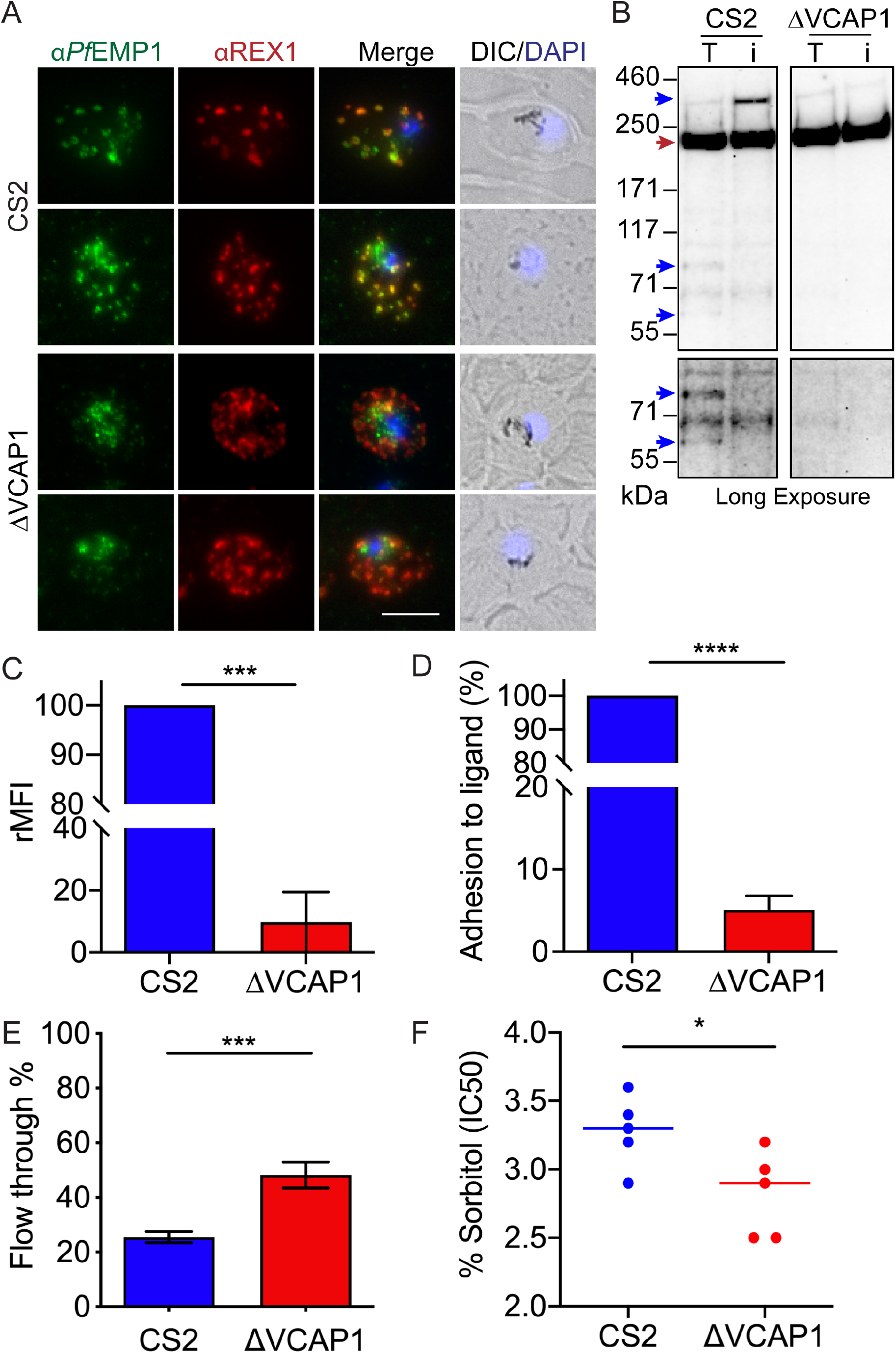
Genetic knockout of VCAP1 lead to a loss of *Pf*EMP1 at the RBC surface and impaired adhesion. (A) Wildtype CS2 and VCAP1 knockout-infected RBCs were fixed and probed with anti-*Pf*EMP1 (green) and counterstained with anti-REX1 (red). Nuclei were stained with DAPI (blue). CS2 = parent line. Projections of Z stacks are shown. Scale bar = 5 μm. (B) Whole infected RBCs were treated with either trypsin (T), or trypsin + trypsin inhibitor (i), solubilised in Triton X-100 and the resultant pellet representing RBC membrane embedded *Pf*EMP1 subjected to western blotting and probed with anti-ATS. Full length *Pf*EMP1 (350 kDa) and ATS bands are indicated with blue arrows. Bands at ∼240 kDa represent cross reaction with spectrin (red arrow). A ∼70 kD cross-reactive species can be seen in all samples. (C) Flow cytometry analysis of infected RBCs labelled with antibodies to the external domains of *Pf*EMP1. Experiments were performed in triplicate on 3 separate occasions. Errors bars represent SEM, Student’s t-test, P = 0.0002. The relative mean fluorescence intensity (rMFI) normalised to CS2 controls is shown. (D) Mid-trophozoite stage-infected RBCs were examined for their ability to bind to chondroitin sulfate-A under physiological flow conditions. The average number of adherent infected-RBCs in 10 fields of view was assessed in 3 independent experiments. Values were normalized to the parent line, CS2. (F) Spleen mimic filtration showing a significant increasing in filterability for the ΔVCAP1 infected RBCs. Experiments were performed in triplicate on 3 separate occasions. Student’s t-test, P = 0.0006. Error bars represent SEM. (G) Analysis of susceptibility to lysis in sorbitol solutions. The IC50 values for triplicate experiments performed on 5 separate occasions are shown. The line represents the median.

We next examined if *Pf*EMP1 is displayed at the surface of ΔVCAP1 infected RBCs by using a trypsin cleavage assay (8, 32). In this assay whole infected RBCs are treated with trypsin (T) or with trypsin plus inhibitor (i), before Triton X-100 solubilisation. Only the Triton-insoluble (RBC membrane-embedded) *Pf*EMP1 population survives the extraction process. In the protease-inhibited CS2 control (i), RBC membrane-embedded full-length *Pf*EMP1 is evident, with an apparent molecular mass of ∼270 kDa (Fig 6B, top blue arrow). It is evident that the pool of RBC membrane-embedded *Pf*EMP1 is decreased in the ΔVCAP1 infected RBCs (Fig 6B, top blue arrow). In the trypsin-treated CS2 sample (T) the surface pool of *Pf*EMP1 is depleted and two cleavage products of ∼75 kDa and ∼60 kDa (blue arrows) which correspond to the membrane embedded conserved ATS region of *Pf*EMP1 were observed. These cleavage products are absent in the ΔVCAP1 infected RBC samples consistent with a loss of *Pf*EMP1 at the RBC membrane (Fig 6B). A cross-reaction of the *Pf*EMP1 antibody with RBC spectrin at ∼220 kDa is observed as well as a number of additional bands of variable intensity which likely represent spectrin breakdown products were present in all samples (21).

To further examine surface presentation of *Pf*EMP1, we used a rabbit polyclonal serum that recognises VAR2CSA on the surface of CS2 parasite-infected RBCs (41). A strong signal was observed in CS2 parasites, which was reduced by 90% in the ΔVCAP1 infected RBCs (Fig 6C). The CS2 parent line expresses a fixed *Pf*EMP1 variant, VAR2CSA, which binds to chondroitin sulphate-A. We investigated infected RBC binding to immobilised chondroitin sulphate-A under physiologically relevant flow conditions. RBCs infected with ΔVCAP1 parasites show a 95% lower level of adhesion compared to the CS2 parent (Fig 6D). This provides further evidence that in the absence of VCAP1, parasites are unable to traffic *Pf*EMP1 to the RBC surface.

### RBCs infected with ΔVCAP1 parasites have increased deformability and fragility

We examined the ability of CS2 and ΔVCAP1 infected RBCs to filter through a bed of beads designed to mimic splenic fenestrations. RBCs infected with ΔVCAP1 parasites synchronised to 20-24 h post invasion (the window during which knob assembly and host cell remodelling occurs (2)) exhibited a 23% increase in the filterability of the parasites compared to CS2 controls (Fig 6E). We next examined the sensitivity of infected RBCs to osmotic lysis. We found that ΔVCAP1 infected RBCs are more sensitive to sorbitol-induced swelling than RBCs infected with wildtype CS2 (Fig 6F).

## Discussion

The suggestion that Maurer’s clefts play a role in transporting virulence proteins from the parasite to the RBC membrane was first put forward nearly 40 years ago (42), however our understanding of the repertoire of cleft-associated proteins and their functions remains incomplete. A previous study examined the proteins present in the ghost fraction of trophozoite stage-infected RBCs and characterised two novel Maurer’s clefts proteins (14). More recently, proteomics approaches have been used to define the “exportomes” of both *P. yoelii* and *P. berghei* (43, 44). In this study, we built on this work and defined the composition of Maurer’s cleft in *P. falciparum* during the late ring stage of infection (14 – 18 h post-invasion). During this time window *Pf*EMP1 is trafficked to Maurer’s clefts and a sub-population is transferred to the RBC membrane (8). A technical advantage of this time window is that the Maurer’s clefts are still mobile in the RBC cytoplasm, allowing separation from host and other parasite components.

Making use of a previously generated REX1-GFP tagged line (17), we used GFP-Trap to affinity purify clefts released by hypotonic lysis. Mass spectrometry analysis identified enriched peptides from 18 proteins. Of these, 11 were previously shown to be Maurer’s cleft-located proteins, and 7 were potentially novel residents.

We also observed enrichment of host cell proteins in the Maurer’s clefts. The most highly enriched protein was Annexin A4, which is involved in vesicle aggregation and the formation of lipid rafts (45). Other enriched human proteins include Annexin A11, which binds to the calcium sensor, calcyclin (33, 46) and Copine-3 (a calcium-dependent phospholipid-binding protein), as well as VPS28, TSG101 (a component of the ESCRT-I complex), syntaxin-7 (a SNARE component) and VAC14 (a scaffold protein involved in phosphatidylinositol 3,5-bisphosphate synthesis). All of these proteins are involved in membrane trafficking. It is possible these proteins play roles in Maurer’s cleft sculpting, vesicle formation or *Pf*EMP1 trafficking.

We generated transfectants expressing GFP fusions of five putative Maurer’s cleft proteins and two known resident proteins. We confirmed that all of the tagged proteins were located at the Maurer’s clefts. The cell lines were then employed in an investigation of protein-protein interactions using GFP-Trap IP and protein identification by mass spectrometry. This analysis identified a network of interactions. We identified hubs where newly identified or poorly characterised Maurer’s clefts proteins participate in networks with known *Pf*EMP1 trafficking proteins and importantly *Pf*EMP1 itself.

For example, we identified a network involving *Pf*EMP1, REX1, PTP5 and the tether protein MAHRP2. This is consistent with previous evidence that PTP5 co-precipitates with GFP-tagged *Pf*EMP1 and that REX1 and PTP5 are needed for *Pf*EMP1 trafficking to the RBC surface (21, 31, 37). We confirmed the interaction between REX1 and *Pf*EMP1, consistent with a previous study showing that GFP-tagged *Pf*EMP1 co-precipitates REX1 and vice versa (31). Our previous work showed that REX1 plays a role in trafficking *Pf*EMP1 from the clefts to the RBC membrane. Using 3D-SIM microscopy we previously showed that REX1 is located at the edges of the Maurer’s clefts (11, 36, 37). In this work, we showed that PTP5 also locates at the cleft periphery, partially overlapping or intercalated with REX1 and *Pf*EMP1. Collectively, these data points to a protein hub that plays a role in the trafficking of *Pf*EMP1 from the Maurer’s clefts to the RBC membrane. Interestingly two members of this peripherally located hub, REX1 and *PF*3D7_1353100, co-precipitate MAHRP2. MAHRP2 is located on parasite-derived structure called tethers that have been proposed to link Maurer’s clefts to the RBC membrane (13).

A second cluster comprises MAHRP1, VCAP1, GEXP10 and PIESP2. 3D-SIM microscopy revealed that MAHRP1, VCAP1 and GEXP10 have an overlapping physical location in the central region of the Maurer’s clefts. Previous work has shown that MAHRP1 disruption prevents trafficking of *Pf*EMP1 to the Maurer’s clefts (35), suggesting a possible role for this protein hub in loading *Pf*EMP1 into the Maurer’s clefts.

We further characterised two proteins from this cluster, VCAP1 and GEXP10. These proteins show sequence similarity (32% identity) and are the only members of the Hyp8 gene family of exported Plasmodium proteins (47). They both contain a signal peptide and a PEXEL motif and have no orthologues in other Plasmodium species. The two proteins were initially reported to be exported to the RBC cytosol in gametocytes (27), but another report identified VCAP1 as a Maurer’s cleft-associated protein in asexual stages (28). Yet another antibody-based study reported that VCAP1 and GEXP10 are located at the RBC surface in the asexual blood stage (29) and are implicated in binding of infected RBCs to the chemokine CX3CL1. That study referred to them as CX3CL1-Binding Proteins 2 and −1 (CBP2 and CBP1, respectively). In our work, we show that both VCAP1 and GEXP10 are present at the Maurer’s cleft with no detectable GFP tagged protein in the RBC cytosol or at the RBC membrane. We therefore classify VCAP1 and GEXP10 as Maurer’s cleft resident proteins but it remains possible they also occupy other locations at levels below the limit of detection in our analysis.

Previous attempts to genetically disrupt VCAP1 and GEXP10 were unsuccessful, suggesting these may be essential (21, 28). Using CRISPR-Cas9 and homology directed repair we were able to successfully generate a ΔVCAP1 parasite line but were unable to generate a GEXP10 knockout. The inability to knock-out GEXP10 is consistent with data from the recent saturation-level mutagenesis study (38) which showed that GEXP10 is indispensable, based on the lack of *piggyBac* transposon insertion sites (Mean Insertion Score of 0.26).

The ΔVCAP1 parasite line exhibited significant ultrastructural changes, with cleft fragmentation and swelling. VCAP1 has two predicted transmembrane domains, as for the previously characterised 2TM-MC Maurer’s cleft protein family. The 2TM-MC proteins have been reported to be oriented with the N- and C-terminal regions facing the RBC cytoplasm (48). If VCAP1 adopts an equivalent orientation, its 26-amino acid loop (between the two transmembrane segments) would face the Maurer’s cleft lumen. It is possible that this loop sequence is involved in interactions that stabilise the Maurer’s clefts, and bring the two lamellae close together. Knock-out of VCAP1 also leads to the formation of enlarged aberrant knobs and knob clusters, sometimes located at the ends of membrane protrusions. Similarly, SEM imaging of the cytoplasmic face of infected RBC membranes reveals doughnut-shaped knob structures that are much larger in the ΔVCAP1 infected RBCs than in the CS2 parent.

The aberrant knob and Maurer’s cleft morphology is associated with a defect in the delivery of *Pf*EMP1 to the RBC membrane, as confirmed by trypsin cleavage assays and flow cytometry using antibodies to the ectodomain of *Pf*EMP1. As a consequence, the binding of the infected RBCs to chondroitin sulphate-A was completely abrogated.

It is interesting to consider how the loss of a Maurer’s cleft-resident protein could result in aberrant knob morphology. While an earlier study using a chimeric KAHRP-GFP fusion (with a non-natural signal sequence) suggested that KAHRP might be trafficked to the knobs via the Maurer’s clefts (49), more recent studies do not support this model. Thus, it seems unlikely that Maurer’s cleft-located VCAP1 plays a direct role in KAHRP trafficking. We considered the possibility that there is a small population of VCAP1 located at the RBC membrane, as previously proposed (29) which may participate in knob assembly. However, we were unable to detect GFP-tagged VCAP1 at the RBC membrane or in the RBC cytoplasm.

An alternative possibility is that VCAP1 is required for the trafficking of an unknown protein required for knob assembly. A recent ultrastructural study visualised the knob complex as a spiral structure connected by multiple links to the RBC membrane skeleton and coated by an electron-dense layer that likely represents KAHRP (50). An earlier cryo-electron tomography study revealed branched actin filaments (each ∼500 nm in length) that connect the Maurer’s clefts to the RBC membrane skeleton in the region of the knobs (12). Recent work from our laboratory suggests that knobs are generated by the trafficking of individual KAHRP-containing modules to the RBC membrane skeleton followed by assembly of KAHRP modules into ring-shaped complexes that sit at the base of the physical knob structure (2). The composition of the spiral core remains unknown. One possibility is that correctly formed Maurer’s clefts may be needed for delivery of a knob assembly factor.

The ΔVCAP1 parasite line exhibited a growth advantage compared to wildtype CS2, perhaps consistent with the general observation that knob-minus parasites outgrow knob-plus parasites in culture. The abnormal knob morphology is associated with increased deformability, suggesting that the reduced number of these aberrant knobs is unable to contribute the strain hardening effects associated with wildtype knobs (51). Of interest, we found that ΔVCAP1 infected RBCs are more sensitive to osmotic changes, suggesting that the abnormal knobs weaken the membrane structure. The current model for *Pf*EMP1 trafficking suggests that it is trafficked across the RBC cytoplasm as a chaperoned complex and then loaded into the membrane at the Maurer’s clefts (52). Forward trafficking delivers *Pf*EMP1 into the RBC membrane bilayer from where it moves laterally to the knobs to form the mature virulence complex (2). The exact mechanism of *Pf*EMP1 unloading and trafficking is not known but it has been speculated that *Pf*EMP1-loaded vesicles bud from the Maurer’s clefts and traffic along remodelled actin filaments or tethers. In the ΔVCAP1 parasite infected RBCs we observe a dramatic fragmentation of the Maurer’s clefts. We propose that this compromises correct trafficking of both PfEMP1 and a knob assembly factor. VCAP1 is a basic protein located in the central region of the Maurer’s clefts. It is possible that VCAP1 interacts with the acidic C-terminal segment of *Pf*EMP1 and facilitates loading into the Maurer’s cleft membrane. It may then transfer *Pf*EMP1 to REX1 (at the Maurer’s cleft periphery) for onward delivery. Further studies are required to fully elucidate the role of VCAP1 in virulence protein trafficking.

In summary, we have developed a novel method for enriching Maurer’s clefts from late ring stage *P. falciparum*-infected RBCs that avoids host cell membrane contamination. We identified a number of novel Maurer’s cleft proteins that expand the repertoire of resident proteins locating at these structures. We characterised different protein hubs that may function in the trafficking of *Pf*EMP1 and determined their spatial organisation. In particular, we identified interacting proteins located in the central region of the Maurer’s clefts that may be involved in loading *Pf*EMP1 into the Maurer’s cleft membrane. In the absence of one of these proteins, VCAP1, the Maurer’s clefts swell and fragment, the knobs enlarge and clump and *Pf*EMP1 trafficking to the RBC membrane fails. RBCs infected with VCAP1-disrupted parasites are more deformable, more sensitive to sorbitol lysis and the parasites possess a growth advantage. A more detailed understanding of the steps in the export pathway may reveal new strategies to target the parasite by blocking the delivery of *Pf*EMP1 to the infected RBC surface.

## Materials and Methods

### Plasmodium falciparum culture

Parasites were cultured as previously described (53). Briefly, *P. falciparum* cell lines were cultured in 5% human O+ red blood cells (Australian Red Cross blood service) in RPMI-GlutaMAX-HEPES (Invitrogen) supplemented with 5% (v/v) human serum (Australian Red Cross blood service), 0.25% (w/v) AlbuMAX II (Invitrogen), 200 μM hypoxanthine, 10 mM D-glucose (Sigma) and 20 μg/ml gentamicin (Sigma). Mature stage parasites were enriched from parasite culture using Percoll purification or magnetic separation (54, 55). Ring stage parasites were enriched by treatment with 5% D-sorbitol (56). Knob positive parasites were enriched by flotation on Gelofusine (Braun) as previously described for gelatin (57). Transfectants containing the pGLUX plasmids were maintained in the presence of 5 nM WR99210 (Jacobus Pharmaceuticals). The ΔVCAP1 parasite line was maintained on 2 nM DSM1 (BEI Resources).

### Plasmid construction and transfection

To create the GFP-tagged transfectant parasites used in this study, the full-length genes (minus the stop codon) were amplified from 3D7 gDNA using primers incorporating *Xho*I and *Kpn*I restriction enzyme cleavage sites at the 5’ and 3’ ends of the gene (Table S9). Sequence verified DNA was then directionally cloned into the *Xho*I and *Kpn*I sites of the pGLUX vector.

To generate the VCAP1 knockout cell lines two regions of homology within the 5’ and 3’ of the VCAP1 locus were PCR amplified using the primers as outlined in Table S9. The HR1 was cloned into the pUFTK plasmid containing a yeast dihydroorotate dehydrogenase cassette which confers resistance to DSM1 using the *Avr*II and *Nco*I sites and HR2 was cloned into the *Spe*I and *Sac*II site of the plasmid. The guide RNA was selected using CHOP CHOP and cloned into the *Btg*ZI linearised pAIO-Cas9 vector containing a hDHFR selection cassette (58). Transfections were performed as previously described (59).

### Live cell and immunofluorescence microscopy

Infected red blood cell smears were fixed in an acetone and methanol solution at −20°C for 10 min. Wells were marked with a hydrophobic PAP pen, then washed three times with PBS. Primary antibodies were diluted in 3% (w/v) bovine serum albumin (BSA) and were added to the wells for 1 h at room temperature and subsequently washed three times with PBS. The primary antibodies used in this study were: mouse anti-REX1 (1:1000, (20)), rabbit anti-REX1 (1:1000, (20)), mouse anti-GFP (1:300, Roche), rabbit anti-GFP (1:300, (60)), mouse anti-MAHRP1 (1:300, (23)), mAb anti-ATS (1:100, (19)), mouse anti-SBP1 (1:300, (18)), rabbit anti-HA (1:300, Sigma Aldrich). Secondary anti-mouse or anti-rabbit AlexaFluor® 488, 568 or 647 antibodies (1:300 in 3% (w/v) BSA, Life Technologies) were added to the wells for 1 h at room temperature, then wells were washed 3 times with PBS. Each well was then treated with DAPI (4’,6-diamidino-2-phenylindole) and PPD antifade (p-phenylenediamine) prior to mounting and sealing. For live-cell fluorescence microscopy, parasites were suspended in RPMI and mounted on a coverslip. Samples were imaged on a DeltaVision Elite Restorative Widefield Deconvolution Imaging System (GE Healthcare). Samples were excited with solid state illumination (Insight SSI, Lumencor). The following filter sets with excitation and emission wavelengths were used: DAPI Ex390/18, Em435/48; FITC, Ex475/28, Em523/26; TRITC, Ex542/27, Em594/45; Cy5 Ex 632/22, 676/34 nm. A 100x UPLS Apo (Olympus, 1.4NA) oil immersion objective lens was used for imaging. For 3D Structured Illumination microscopy (3D-SIM), the DeltaVision OMX V4 Blaze was used (GE Healthcare). Samples were imaged using a 60X Olympus Plan APO N (1.42 NA) oil immersion lens. The following laser emission and band pass filter sets were used: Ex488, Em528/48; Ex568, Em609/37 or Ex642, Em683/40 nm. Images were processed using the FIJI ImageJ software (61).

### SDS-PAGE and immunoblotting of protein samples

For immunoblotting, samples were resuspended in Bolt™ LDS sample buffer and 50 mM dithiothreitol (DTT) and separated on Bolt™ 4-12% Bis-Tris gel at 200V for 32 min. Separated samples were transferred to a nitrocellulose membrane using an iBlot® gel transfer device. Membranes were blocked in 3% (w/v) skim milk for 1 h at room temperature. The membranes were then probed for 16 h at 4 °C with the following primary antibodies in 3% (w/v) skim milk in PBS: mouse anti-GFP (1:1000, Roche), rabbit anti-SBP1 (1:1000, (18)), mAb anti-ATS (1:100, (19)), mouse anti-REX1 (1:1000, (20)), mouse anti-EXP1 (1:1000), rabbit anti-spectrin (1:1000, Sigma Aldrich), mouse anti-MAHRP1 (1:1000, (23)), rabbit anti-GAPDH (1:1000), rabbit anti-HA (1:1000, Sigma Aldrich). Membranes were washed 3 times in 0.05% (v/v) Tween-20 in PBS for 10 min. Secondary antibodies (anti-mouse or anti-rabbit) conjugated to horseradish peroxidase (HRP, Promega) were diluted 1:25,000 in 3% (w/v) skim milk in PBS and were incubated with membranes for 1 h at room temperature. Membranes were again washed 3 times in 0.05% (v/v) Tween-20 in PBS, then once in PBS before being incubated with Clarity™ ECL reagents (Bio-Rad) and visualised on a ChemiDoc imaging system (Bio-Rad).

### Enrichment of Maurer’s clefts

Parasites were synchronised to a 4 h window by Percoll purification followed by sorbitol lysis. At 14-18 h post-invasion, parasites (∼15% parasitaemia) were harvested and washed in PBS. Infected red blood cells were lysed on ice with chilled hypotonic buffer (1 mM HEPES/NaOH, Roche cOmplete EDTA-free protease inhibitor, pH 7.4). The lysed infected red blood cell solution was passaged through a 27-gauge needle 10 times and was then made isotonic by the addition of 4X assay buffer (200 mM Hepes-NaOH, 200mM NaCl, 8 mM EDTA (pH 7.4). The solution was centrifuged at 2500 g for 10 min at 4 °C, the supernatant was collected and centrifuged again at 2500 g for 10 min at 4 °C. The supernatant (containing the Maurer’s clefts) was then pre-cleared with Pierce™ Protein A agarose beads (Thermo Scientific) for 30 min at 4 °C on a mixing wheel. The Protein A agarose beads were pelleted by centrifugation and the supernatant was incubated with GFP-Trap® agarose beads (Chromotek) for 4 h at 4 °C on a mixing wheel. The beads were then collected and used for downstream applications.

### Co-IP with GFP-Trap agarose beads

Parasite-infected RBCs were purified using a VarioMACS™ (Miltenyi Biotec) magnet or by floatation in 70% (v/v) Gelofusine® (Braun) in PBS. The infected RBCs were washed in RPMI and then solubilised on ice for 30 min in 10 times pellet volumes of IP buffer (1% (v/v) Triton X-100, 150 mM NaCl, 50 mM Tris, 8 mM EDTA) with Complete™ protease inhibitor cocktail (Roche). The Triton X-100-insoluble material was pelleted twice by centrifugation (10 min at 16,000 *g*). The supernatant was pre-cleared with Pierce™ Protein A agarose beads (Thermo Scientific) for 1 h at 4 °C on a mixing wheel. The Protein A beads were pelleted and an aliquot of the supernatant was taken as the input fraction. The remainder of the supernatant was incubated with washed GFP-Trap agarose beads (Chromotek) for 16 h at 4 °C. The GFP-Trap beads were then washed 5 times in IP buffer.

### Mass spectrometric analysis of Maurer’s clefts and co-IPs

Agarose beads were washed twice in 1 mM Tris-HCl prior to elution of bound proteins. Samples were prepared for mass spectrometric analysis as previously described (31). Briefly, proteins were eluted from GFP-Trap beads by the addition of 20% (v/v) trifluoroethanol in formic acid (0.1%, pH 2.4) and were incubated at 50°C for 5 min. The eluate was reduced with 5 mM TCEP (Thermo Fischer Scientific) and neutralised with TEAB (tetraethylammonium bromide). Samples were digested with trypsin (Sigma) overnight at 37 °C.

Samples were analysed by ESI LC-MS/MS on Orbitrap Elite (purified Maurer’s clefts) or Q Exactive (co-IPs) mass spectrometers. Mass spectra (ProteoWizard) were searched against a custom database containing the *Plasmodium falciparum* 3D7 and UniProt human proteomes. Searches were performed on MASCOT (Matrix Science), with the following parameters: precursor ion mass tolerance of 10 ppm, fragment ion mass tolerance of 0.2 Da, trypsin as the cleavage enzyme, three allowed missed cleavages and allowing for oxidation. The network map of protein interactions was created using NAViGaTOR 2.3 software, networks were redrawn in Adobe Illustrator. The mass spectrometry proteomics data have been deposited to the ProteomeXchange Consortium via the PRIDE (62) partner repository with the dataset identifier PXD014873. (Tempory Username: Password: thU0KxPW).

### Infected RBC binding assay under flow conditions

Ibidi µ-Slide 0.2 channel slides were incubated with 100 µl chondroitin sulfate A (100 µg/ml, Sigma) in 1% (w/v) BSA in PBS overnight at 37°C. Channels were blocked with 1% (w/v) BSA in PBS for 1 h at room temperature then gently flushed with warm bicarbonate-free RPMI 1640 (Invitrogen). Mid-trophozoite stage cultures were diluted to 3% parasitemia and 1% haematocrit in bicarbonate-free RPMI 1640 and pulled through the channel at 100 µl/min for 10 min at 37°C. Unbound cells were washed out of the channel at 100 µl/min for 10 min at 37°C. Adherent cells were counted at 10 points along the axis of the channel.

### VAR2CSA ectodomain labelling and analysis via flow cytometry

Mid-trophozoite cultures were diluted to 3% parasitemia, 0.4% hematocrit and plated into a 96 well plate. Cells were washed 3 times in 1% (w/v) BSA in PBS and incubated with a rabbit polyclonal anti-VAR2CSA for 30 min at 37°C (R1945 1:50, Elliott et *al.*, 2005) or a 1% (w/v) BSA in PBS control. Cells were washed in 1% BSA/PBS between all subsequent incubations and all antibodies were diluted in 1% BSA/PBS. Cells were incubated with mouse anti-rabbit IgG (Dako; 1:100) for 30 min at 37°C, then mouse anti-GFP (Alexa Fluor-488) under the same conditions. Following tertiary labelling, cells were washed 3 times in 1% (w/v) BSA in PBS. Nuclei were stained with SYTO61 as described previously (Klonis et *al.*, 2011) and analysed via flow cytometer (BD FACSCanto II). SYTO61 positive cells were gated and the Alexa Fluor-488 fluorescence intensity measured. The results are displayed as the percentage of fluorescence relative to wild type controls.

### Trypsin cleavage of surface exposed *Pf*EMP1

Late stage infected RBCs were purified and incubated with 20 volumes of 1 mg/ml TPCK-treated trypsin in PBS (Sigma) or trypsin and soybean trypsin inhibitor (5 mg/ml, Sigma) at 37°C for 1 h. Trypsin activity was quenched with a 15 min incubation with trypsin inhibitor at room temperature for 15 min. Cells were subsequently solubilised in 1% Triton X-100 in PBS containing 1x cOmplete protease inhibitor (Roche) and incubated on ice for 20 min. The sample was centrifuged at 16,000 x g at 4C for 10 min and the resulting pellet washed in the triton solution 2 additional times. The resulting triton-insoluble pellet was solubilized in 2% SDS in PBS and mixed on a rotating wheel for 40 min. This SDS-soluble fraction was then separated by SDS-PAGE and transferred to a nitrocellulose membrane for western blotting.

### Parasite growth assay

Parental CS2 and ΔVCAP1 cell lines were synchronised to a 2 h window and diluted to 1.5% late stage parasitemia. Every 48 h infected red blood cells were taken from culture and stained with SYTO-61 nuclear stain and parasitemia was calculated by flow cytometry as previously described (Klonis et *al.*, 2011). The cultures were then diluted by the same dilution factor. Parasitaemia was recorded over 4 cycles and normalised to the parental CS2 line.

### Sorbitol sensitivity assay

Mid-trophozoite infected red blood cells at 2-5% parasitemia were incubated at 37°C with one of 12 solutions between 0.1% (w/v) and 5.5% (w/v) D-sorbitol. Parasitemia was measured via flow cytometry following incubation with nuclear stain SYTO-61 and 30,000-50,000 events recorded per well as previously described (Klonis et *al.*, 2011). Parasitemia was normalised to an internal RPMI control for each cell. This data was then used to calculate the IC50 values for each separate experiment. The IC50 values from each experiment are plotted.

### Microbead filtration

Spleen mimic filtration was performed to assess the deformability properties of parasite infected RBC (63). Synchronous parasites at 22 h post-invasion were prepared at 6% parasitemia and 1% hematocrit in 1% (w/v) AlbuMAX II in 1X PBS and flowed through the microbead bed using a syringe pump at a flow rate of 60mL/min. The data presented is the percentage of parasites present in the flow through relative to starting parasitemia. The mean and standard error are plotted, from 3 separate experiments. A Students t-test was used to evaluate statistical significance.

### Membrane shearing

Glass coverslips were cleaned with acetone and 50% methanol prior to treatment with 3-aminopropyl triethoxysilane (APTES), bis-sulfosuccimidyl suberate and addition of the ligand erythroagglutinating phytohemagglutinin (PHAE) as previously described (64). Infected RBCs were immobilised on the functionalised glass slides, then sheared by applying a hypotonic buffer (5 mM Na_2_HPO_4_/NaH_2_PO_4_, 10 mM NaCl, pH 8) from a 30 mL syringe (23-G needle) at an angle of ∼20° (65). The membrane disks were placed in PBS prior to down-stream imaging.

### Scanning Electron Microscopy

Whole infected RBC cell samples bound to glass coverslips were immersed in 0.05% glutaraldehyde in PBS for 20 minutes prior to fixation in 2.5% glutaraldehyde for 2 h at room temperature. Sheared membranes were prepared for scanning electron microscopy as previously described (2). In brief, sheared cells were fixed immediately after lysis with 2.5% glutaraldehyde in PBS for 2 h at room temperature. Both whole and sheared cell samples were then washed thoroughly with deionised water and dehydrated via sequential 5 min incubations in 20%, 50%, 70%, 80%, 90%, 95% and (3x) 100% ethanol before critical point drying in a Leica CPD300 critical point dryer. Samples were stored under desiccation and gold coated immediately before imaging. Coverslips were gold coated for 40 s or 75 s at 25 mA using a Dynavac sputter coating instrument to a thickness of ∼0.2 nm or ∼0.4 nm for sheared and whole cells respectively. Images were acquired with the ETD detector of an FEI Teneo SEM in Optiplan mode, at a working distance of 5 mm, a beam current of 50 pA and a 2-kV accelerating voltage.

### Immuno-electron Tomography

Infected red blood cells (20 – 30 h post invasion) were magnet purified, washed in PBS, and fixed in 10 pellet volumes of 2% (v/v) *PF*A in PBS for 20 min at room temperature. Cells were washed then permeabilised in 10 pellet volumes of PBS with 1 HU equinatoxin II for 6 min. After washing cells were fixed again in 2% PFA/PBS for 5 min, washed, then blocked in suspension for 1 h with 3% BSA/PBS. Cells were incubated with anti-REX1 (1:50) for 2 h, washed, then incubated with a gold secondary for 1 h (1:15, Aurion protein A EM grade 6 nm gold JA806-111). Cells were washed in 3% (w/v) BSA in PBS then in PBS to remove the BSA.

Cell pellets were resuspended and fixed in 2.5% glutaraldehyde at 4°C for at least 1 h, prior to pre-embedding in low melting point agarose. Cells were then post fixed in 1% potassium ferricyanide (K_3_[Fe(CN)_6_]) reduced osmium tetroxide (OsO_4_) solution for 1 h. Blocks were then rinsed 5 x 3 min in ddH_2_O then dehydrated by sequential incubation 5 min each in 20%, 50%, 70%, 80%, 90%, 95%, and 100% ethanol. Blocks were incubated in 100 % ethanol twice more 5 min each, followed by ethanol/acetone solution (1:1) incubation for 30 min, then 100% acetone for 30 min. Acetone was substituted for acetone/1% thiocarbohydrazide resin (1:1) for 2 h and infiltrated with 100% resin 2 x 12 h and polymerised in oven at 60°C.

Resin blocks were trimmed and sectioned on ultramicrotome (Leica EM UC7, Leica Microsystems, Wetzlar, Germany) and both 70 nm and 300 nm sections were prepared for imaging and electron tomography, respectively. The sections were then post stained by 4% uranyl acetate in water for 10 mins and Reynold’s lead citrated for 10 min. The 300 nm sections were overlayed with 10 nm fiducial gold particles on both sides of the grid. Imaging and electron tomography were performed on FEI Tecnai F30 electron microscope (FEI company, Hillsboro, US) at an accelerating voltage of 300kV. The tilt series were acquired for every 2° in the range between −70° to 70° tilts. Virtual sections were reconstructed from raw tilt series in IMOD using weighted back projection algorithm (66).

## Acknowledgements

The authors thank the Australian Red Cross Blood Service. We thank Dianne Taylor for anti-KAHRP antibodies, Stephen Rogerson for VAR2CSA antibodies and Alan Cowman for the anti-ATS antibodies. LT is a Georgina Sweet, Australian Research Council Laureate Fellow (LE150100011) (http://www.arc.gov.au). MWAD and LT thank the Australian Research Council (DP110100624) (http://www.arc.gov.au) and the National Health and Medical Research Council (1098992) (https://www.nhmrc.gov.au) for funding this work. MWAD was supported by a National Health and Medical Research Council Training fellowship (602541). We thank Arman Namvar for technical assistance. Super-resolution microscopy was performed at the Biological Optical Microscopy Platform and electron microscopy at the Bio21 Institute Advanced Microscopy Facility, The University of Melbourne (www.microscopy.unimelb.edu.au). Mass spectrometry was performed at the University of Melbourne Mass Spectrometry and Proteomics Facility.

## Supplementary Figure Legends

**Fig S1. Live-cell microscopy of REX1-GFP bound to GFP-Trap beads and full-length Western blots of GFP-tagged transfectants.**

(A) Fluorescence microscopy of the beads reveals structures with dimensions consistent with a mixture of intact (∼500 nm) and fragmented Maurer’s clefts. Scale bar = 5 μm. (B) Full uncropped Western blot images for assessing enrichment of Maurer’s clefts from REX1-GFP infected RBCs. These blots were cut horizontally in some cases to simultaneously probe for proteins of different sizes. Total = complete infected RBC lysate; Input = fraction applied to GFP-Trap beads; Unbound = supernatant after GFP-Trap bead incubation, Beads = material bound to beads. Molecular masses are in kDa. (C) Western blot of saponin-lysed infected RBCs from GFP-tagged transfectant parasite lines. Green arrowheads indicate predicted size of the mature full-length protein. Molecular masses are in kDa.

**Fig S2. Immunofluorescence microscopy and Western blots of GFP-tagged proteins**

(A) The GFP-tagged transfectant-infected RBC smears fixed in acetone/methanol (9:1) at −20°C. Samples were labelled with anti-GFP (green) to localise the GFP-tagged protein and anti-REX1 (red) to label the Maurer’s clefts. Parasite nuclei were stained with DAPI (blue). Scale bars = 5 μm. (B-E) Infected RBCs were solubilised in 1% Triton X-100 and subjected to SDS-PAGE. Input loading was 4% of the pellet loading. Green arrowheads indicate the full-length GFP-tagged proteins. Blue arrowheads indicate co-precipitated proteins. (B) 3D7 wildtype, REX1-GFP; (C) PTP6-GFP; (D) GEXP10-GFP, VCAP1-GFP, MAHRP1-GFP (M1), *PF*3D7_1353100-GFP (*PF*13) IPs, probed with anti-GFP antibody. (E) GEXP10-GFP, VCAP1-GFP and MAHRP1-GFP (M1) IPs probed with anti MAHRP1 antibodies. Input (In) and IP fractions (IP) are shown. Molecular masses are in kDa.

**Fig S3. Complete network map of co-IP experiments**

(A) Connectivity map of the GFP tagged bait proteins and their interactions with proteins identified in the co-IP experiments. (B) Connectivity map combining IP data for SEMP1, PTP1 and *Pf*EMP1 from the literature. Green nodes are the GFP-tagged proteins. Blue nodes are exported interacting proteins. Purple nodes represent proteins within the parasitophorous vacuole. Double width lines represent reciprocal IP.

**Fig S4. Whole cell analyses of GFP tagged transfectants by 3D-SIM.**

REX1-GFP-infected RBCs were fixed and labelled with anti-GFP (green) and anti-MAHRP1 (red). (B) The remaining GFP-tagged transfectant-infected RBCs were fixed and labelled with anti-GFP (green) and anti-REX (red). Average projections of z-stacks are displayed. The merged image and zoom as displayed in Fig 3 are shown. Merge scale bars = 3 μm. Zoom scale bars = 3 μm. (B) 3D-SIM imaging of GEXP10-GFP and REX1-GFP (red) parasites co-stained with ATS antibodies (*Pf*EMP1) (green). Merged images and a zoom of the merge are shown. Merge scale bars = 3 μm. Zoom scale bars = 1 μm.

**Fig S5. Generation and validation of the ΔVCAP1 parasite line.**

(A) Gene disruption and PCR validation schematic. GOI = gene of interest; HR1 = homology region 1; HR2 = homology region 2; crossed lines indicate crossover event. Arrows, primers; bracketed lines, expected amplicon size. (B) PCR amplicons, letters in reference to primers in part B; KO = knockout; first lane features the marker. (C) Long term growth assay. Cumulative growth rates of wild type CS2 and ΔVCAP1 parasite lines over 4 parasite lifecycles. n=4.

**Fig S6. Whole cell images showing SEM and TEM of CS2 and ΔVCAP1 infected RBCs**

(A) TEM of Equinatoxin treated RBCs highlighting the fragmented Maurer’s clefts. (B) Thin section TEM. EQII permeabilised cells large section view of the zoomed Maurer’s clefts shown in fig 4 C. Parasite features are labelled. K = knobs; MC = Maurer’s Clefts. Scale bars 500 nm.

**Fig S7. Knob morphology examples and Immunofluorescence microscopy of *Pf*EMP1 location in ΔVCAP1 infected RBCs**

(A) Scanning electron microscopy images of normal, large, clustered and asymmetrical knobs. Scale bars = 100 nm. (B) Enhanced exposure images of **Δ**VCAP1 infected RBCs from figure 6A. Anti-*Pf*EMP1 (green) and counterstained with anti-REX1 (red). Nuclei were stained with DAPI (blue). Projections of Z stacks are shown. Scale bar = 5 μm.

**Movie S1. MC mobility movie**

Movie showing the mobility of the Maurer’s clefts (Green) in 14-18 h post-invasion parasite infected RBCs.

**Movie S2. CS2 electron tomography translation and rendered model**

Part 1: Movie translating through a reconstructed z-stack for CS2 infected red blood cells that were equinatoxin II permeabilised and probed with anti-REX1 (1:50) followed by Aurion protein A (EM grade 6 nm gold JA806-111, 1:15). Part 2: 360° rotation around the y axis of CS2 Maurer’s clefts 3D structure reconstructed and rendered in IMOD. Red = RBC; fuchsia stalk = tether; pastel hues = independent clefts; scale bar = 100 nm.

**Movie S3. ΔVCAP1 electron tomography translation and rendered model**

Part 1: Movie translating through a reconstructed z-stack for ΔVCAP1 infected red blood cells that were equinatoxin II permeabilised and probed with anti-REX1 (1:50) followed by Aurion protein A (EM grade 6 nm gold JA806-111, 1:15). Part 2: 360° rotation around the y axis of ΔVCAP1 Maurer’s clefts 3D structure reconstructed and rendered in IMOD. Red = RBC; fuchsia stalk = tether; pastel hues = independent clefts; scale bar = 100 nm.

## Tables

All tables are available for download from figshare using the following link. https://melbourne.figshare.com/s/ab61640da1d1849f0887

## Data sets

All data sets are available for download from figshare using the following link. https://melbourne.figshare.com/s/2cdf8cc5b787d6b8ea67

